# Promoter activity of ORF-less gene cassettes isolated from the oral metagenome

**DOI:** 10.1101/427781

**Authors:** Supathep Tansirichaiya, Peter Mullany, Adam P. Roberts

**Affiliations:** Department of Microbial Diseases, University College London, Eastman Dental Institute, 256 Gray’s Inn Road, London, WC1X 8LD, UK; Liverpool School of Tropical Medicine, Pembroke Place, Liverpool, L3 5QA, UK; Department of Clinical Dentistry, Faculty of Health Sciences, UiT the Arctic University of Norway, Tromsø, Norway

**Keywords:** integron, ORF-less gene cassettes, promoter activity, oral metagenome

## Abstract

Integrons are genetic elements consisting of a functional platform for recombination and expression of gene cassettes (GCs). GCs usually carry promoter-less open reading frames (ORFs), encoding proteins with various functions including antibiotic resistance. The transcription of GCs relies mainly on a cassette promoter (P_C_), located upstream of an array of GCs. Some integron GCs, called ORF-less GCs, contain no identifiable ORF with a small number shown to be involved in antisense mRNA mediated gene regulation.

In this study, promoter sequences were identified, using *in silico* analysis, within GCs PCR amplified from the oral metagenome. The promoter activity of ORF-less GCs was verified by cloning them upstream of a *gusA* reporter, proving they can function as a promoter, presumably allowing bacteria to adapt to multiple stresses within the complex physico-chemical environment of the human oral cavity. A bi-directional promoter detection system was also developed allowing direct identification of clones with promoter-containing GCs on agar plates. Novel promoter-containing GCs were identified from the human oral metagenomic DNA using this construct, called pBiDiPD.

This is the first demonstration and detection of promoter activity of ORF-less GCs and the development of an agar plate-based detection system will enable similar studies in other environments.

## Introduction

Integrons are bacterial genetic elements able to integrate and express genes present on gene cassettes (GCs) [1-3]. They consist of two main components; a functional platform and a variable array of GCs. The functional platform, located on the 5’ end of an integron, consists of an integrase gene (*intI*), and its promoter (P*_intI_*), an *attI* recombination site and a constitutive cassette promoter (P_C_) for the expression of GCs [4]. IntI is a site-specific tyrosine integrase that catalyses the insertion and excision of GCs via recombination mainly at *attI* and the *attC*, the latter located at the joint of excised, circularised GCs. The integrase gene; *intI*, is normally transcribed in the opposite direction to GCs within an integron (Fig 1A). However, some integrons have integrase genes transcribed in the same directions as their GCs. These are called unusual integrons or reverse integrons (Fig 1B), and have been identified in *Treponema denticola, Chlorobium phaeobacteroides* and *Blastopirellula marina* [5, 6].

**Figure 1:**
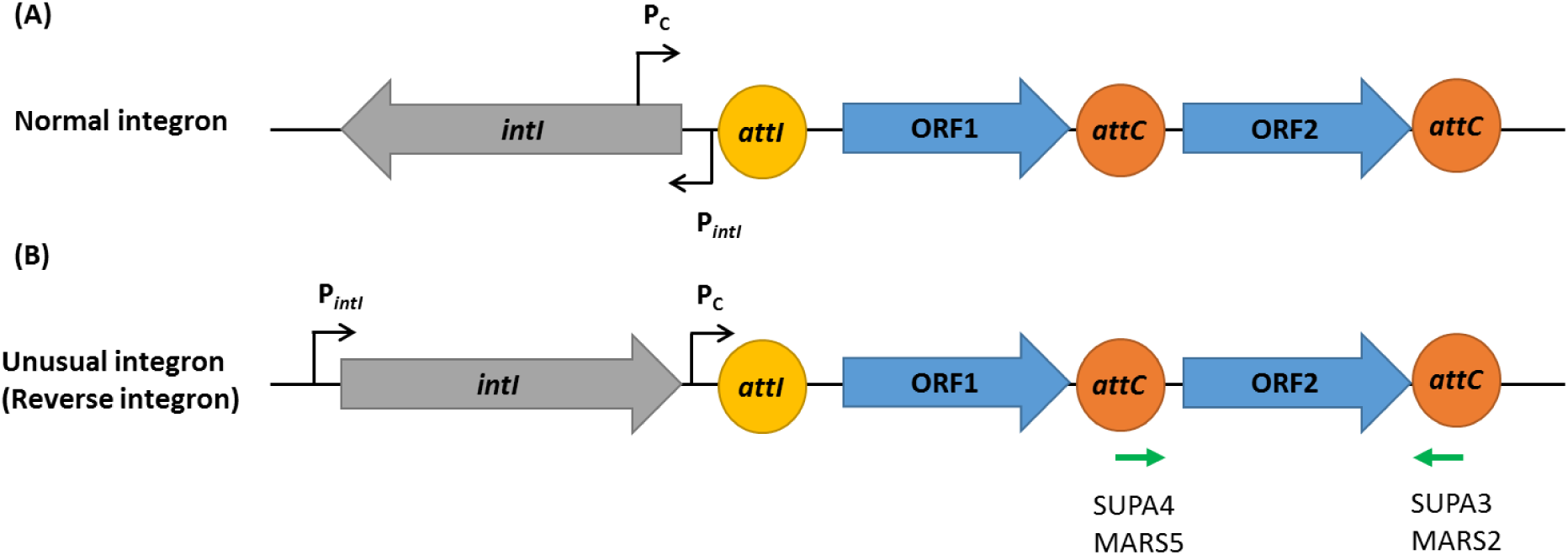
A generalised structure of (A) usual integrons and (B) unusual, or reverse integrons. The green arrows indicate the primer binding sites on the unusual integron structure of *T. denticola*. The grey and blue open arrowed boxes represent integrase gene *(intI)* and the open reading frames (ORFs), respectively, pointing in the direction of transcription. The promoters, *P_intI_* and P_C_, were represented by black arrows. The recombination sites, *attI* and *attC*, were represented by yellow and orange circles, respectively.

The second part of an integron is an array of GCs. Each usually contains a single promoterless open reading frame (ORF) and an *attC* recombination site [7]. The proteins encoded by GCs are diverse and include those associated with antibiotic resistance, virulence, and metabolism [2, 8]. When a GC is excised from integron, it forms a non-replicative mobile genetic element, which can be a substrate for integrase mediated recombination between *attI* (on the integrons) and *attC* (on the circular GC). This directionality of recombination is favoured over *attC:attC* recombination, resulting in the usual insertion of a newly integrated GC immediately next to the P_C_ promoter in the first position of GC array, ensuring maximal expression [9-11].

The expression of integron integrases is controlled via the SOS response; there is a LexA-binding site located in the *P_intI_* [12]. In the absence of stress, the transcriptional repressor LexA binds to *P_intL_* and prevents the transcription of *intI*. The SOS response is activated upon the accumulation of single-strand DNA (ssDNA), generated during DNA damage, DNA repair, transformation, conjugation and certain antibiotic exposure e.g. trimethoprim and fluoroquinolones [13-15]. RecA recognises ssDNA and polymerises into RecA nucleofilaments, which induce autocleavage of LexA, releasing *P_intI_* from repression and allowing *intI* transcription [12, 16]. By controlling the expression of IntI, bacteria can reshuffle their GCs at the precise moments of need (stress), thereby avoiding accumulation of random recombination events that could be deleterious to the host cell [17, 18].

As most of the GCs do not contain a promoter, their expression usually relies on the P_C_ promoter. The level of expression of GCs varies depending on the distance from P_C_, as the strength of expression decreases when GCs are located further from P_C_ [19]. This ensures that a recently acquired GC will be immediately expressed ensuring rapid adaptation due to stress-induced repositioned gene within the integron GC array. There are also some GCs that contain their own promoters, ensuring constitutive expression of their genes regardless of the P_C_ promoter and their position within the integron array; examples include *cmlA1* (chloramphenicol resistance), *qnrVC1* (quinolone resistance), *ere(A)* (erythromycin resistance) and many of the GCs encoding toxin-antitoxin (TA) systems [20-23].

Integron GCs have been identified from environments such as soils, marine sediments, seawater and more recently from human oral metagenomes [24-28]. In our previous study on the detection of integron GCs in the human oral metagenome, we found 13 ORF-less GCs out of 63 identified GCs (20%) [28]. ORF-less GCs have been shown to encode regulatory RNAs, for example the trans-acting small RNA (sRNA)-Xcc1, encoded by the ORF-less GC of a *Xanthomonas campestris* pv. *campestris* integron, which is involved in regulation of virulence [29]. Whilst promoter activity of ORF-less GCs has been discussed, this has not been experimentally demonstrated [8].

In this study, we performed *in silico* analysis to identify promoter sequences in the GCs identified in our previous study on the oral metagenome. Promoter activity was experimentally determined by cloning the selected GCs upstream of the *gusA* reporter gene and measuring β-glucuronidase enzyme activity. Furthermore, we devised a GC-based promoter detection strategy utilising PCR and subsequent cloning between divergently orientated reporter genes. With this system, the successful cloning of amplicons from promoter-containing GCs can be visualised directly on agar plates, allowing the direct isolation of GC PCR amplicons with promoter activity from metagenomic DNA.

## Results

### *in silico* analysis of the promoter sequences on the ORF-less GCs

Among 63 GCs previously identified from human oral metagenomic DNA, 13 were predicted to be ORF-less GCs [28]. Using BPROM promoter prediction software, all ORF-less GCs were predicted to contain promoters on both strands, suggesting that these GCs can transcribe genes in flanking GCs (Table 1). In this study, we have defined the sense strand as the same strand containing the P_C_ promoter (Fig 1).

**Table 1;.**
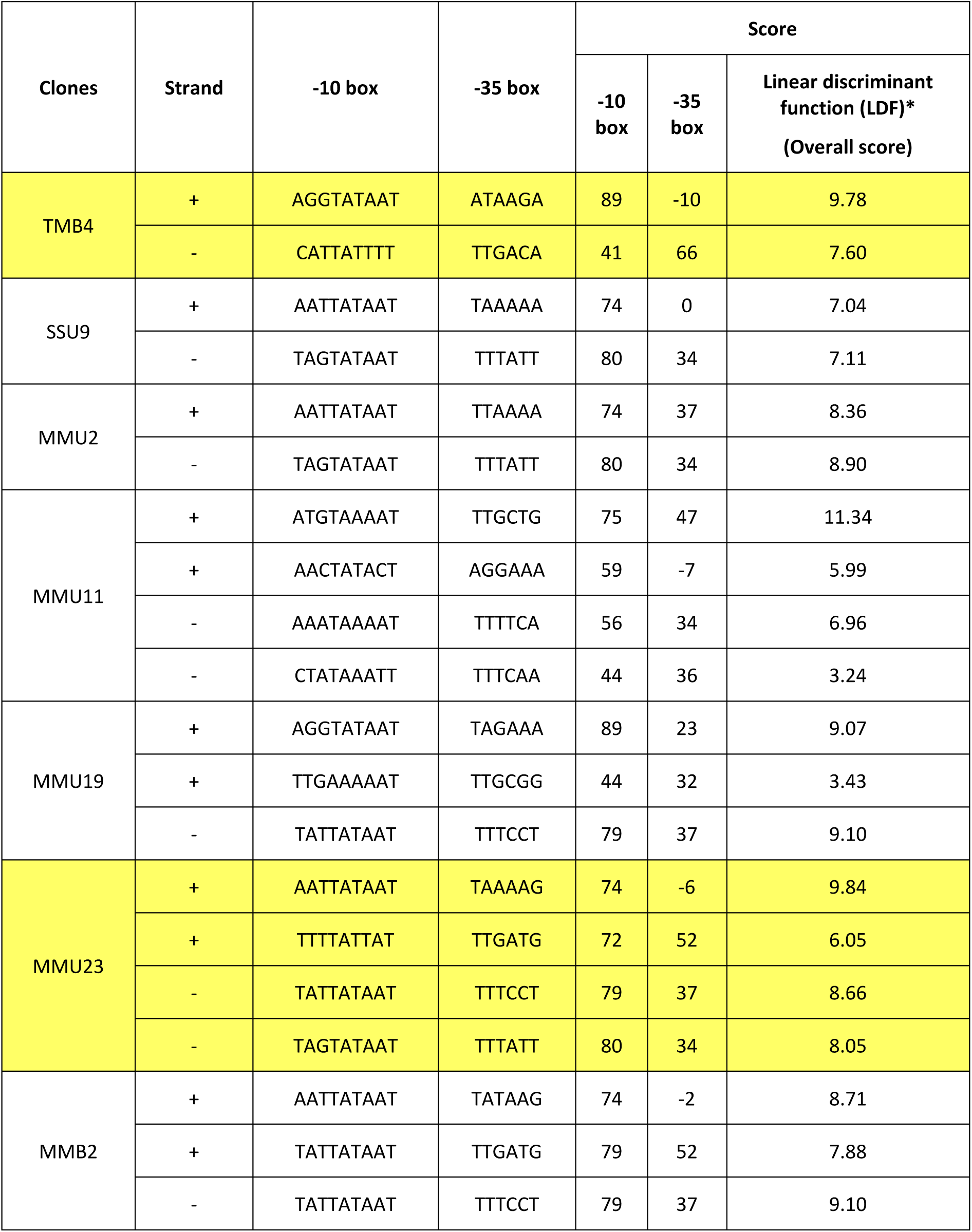
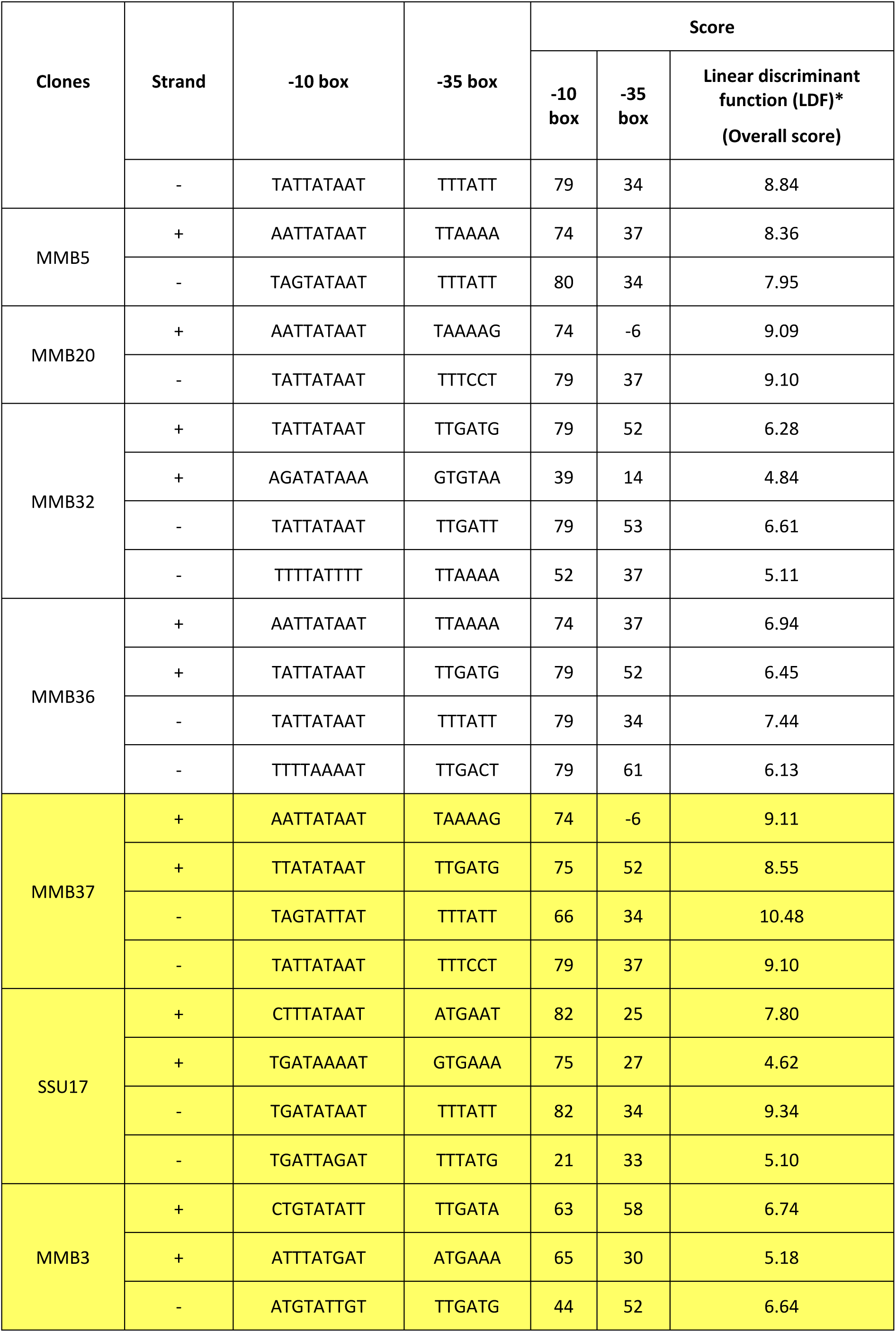
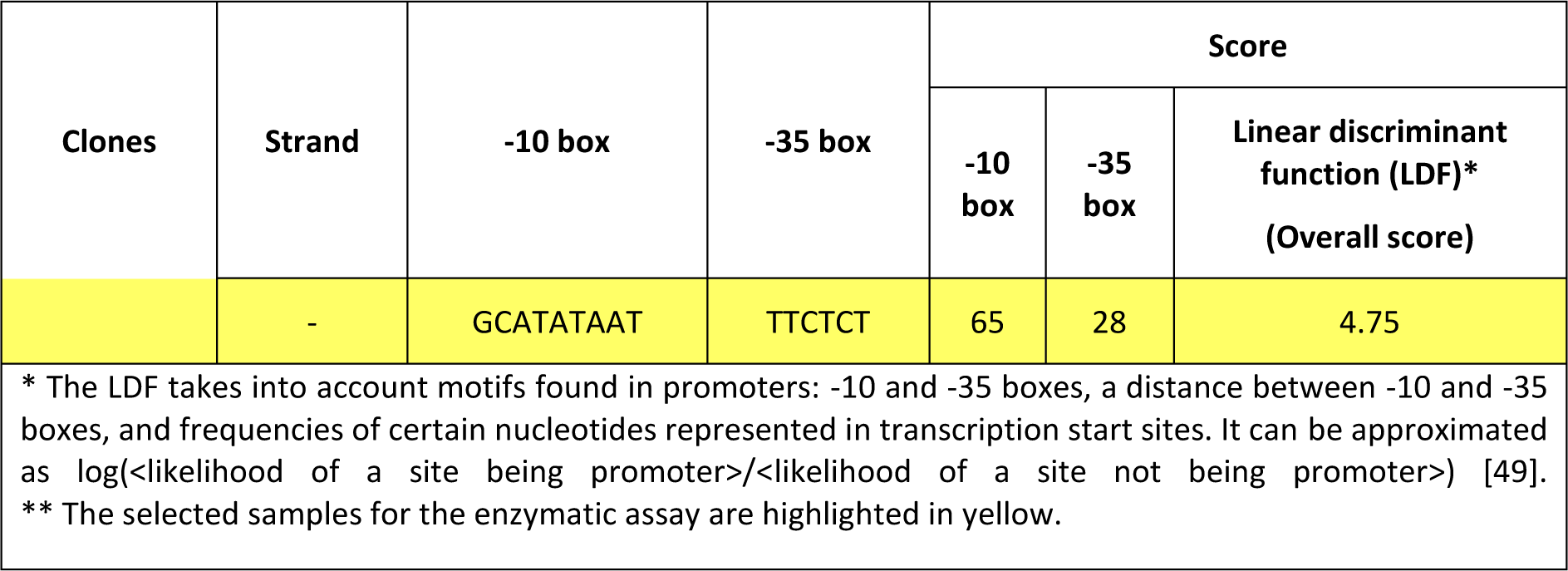
The putative promoters for ORF-less GCs and GCs with an ORF (SSU17 and MMB3) predicted using BPROM.

### Determination of promoter activity of the ORF-less GCs using the β-glucuronidase assay

Five GCs were chosen for experimental expression analysis. GC TMB4 (amplified with primers targeting *intI* and *attC)* was selected as it is ORF-less and located in the first position of the integron array [28]. ORF-less GCs MMU23 and MMB37 were selected as they had the highest overall score predicted by BPROM. Finally, GCs SSU17 and MMB3 were selected as controls, to represent GCs with an ORF.

As BPROM predicted putative promoter sequences on both strands, promoter activity of the selected GCs was determined by directionally cloning upstream of a promoterless β-glucuronidase *(gusA)* gene on *pCC1BAC-lacZα-gusA* (Fig 2) in both directions. For the first position GC; TMB4, three different constructs were made: TMB4 P_C_ promoter, TMB4 GC, and TMB4 P_C_-GC constructs. As the TMB4 P_C_ promoter was not identical to the P_C_ of *T. denticola* integron [30], the P_C_ of another integron GC; TMB1 [28], which was identical to it [30], was included. As the selected GCs were likely derived from *Treponema* spp., two experimentally verified *T. denticola* promoters, PTdTro and PFla, were also included as controls showing that *T. denticola* promoters can be recognised in our *E. coli* host [31, 32]. P_Fla_ and P_Tdtro_ were selected as they rely on different sigma factors. P_Tdtro_ is recognised by sigma factor 70 (σ^70^) that is responsible for the transcription of most genes during growth in both *E. coli* and *Treponema* spp. [31, 33], while P_Fla_ is recognised by sigma factor 28 (σ^28^), involved in the expression of flagella-related genes in motile bacteria [32, 34]. This will determine the limitations of our assay in recognising promoters associated with different types of sigma factors. The results are shown in figure 3. MMB37 and MMB3 had promoter activity on one strand, while MMU23 and SSU17 had no promoter activity, compared to the negative control. The TMB4-P_C_, TMB4 GC, TMB4 P_C_-GC and TMB1-P_C_ constructs, all showed promoter activities on both strands. The P_Tdtro_ from *T. denticola* showed strong promoter activity on both sense and antisense strands, verifying that promoters from *T. denticola* are recognised by *E. coli*. As the P_C_ promoter sequences on TMB1 and TMB4 samples were different at several nucleotides, it was shown that TMB4-P_C_ had higher promoter activities than the TMB1-P_C_ in both directions (Fig 3).

**Figure 2:**
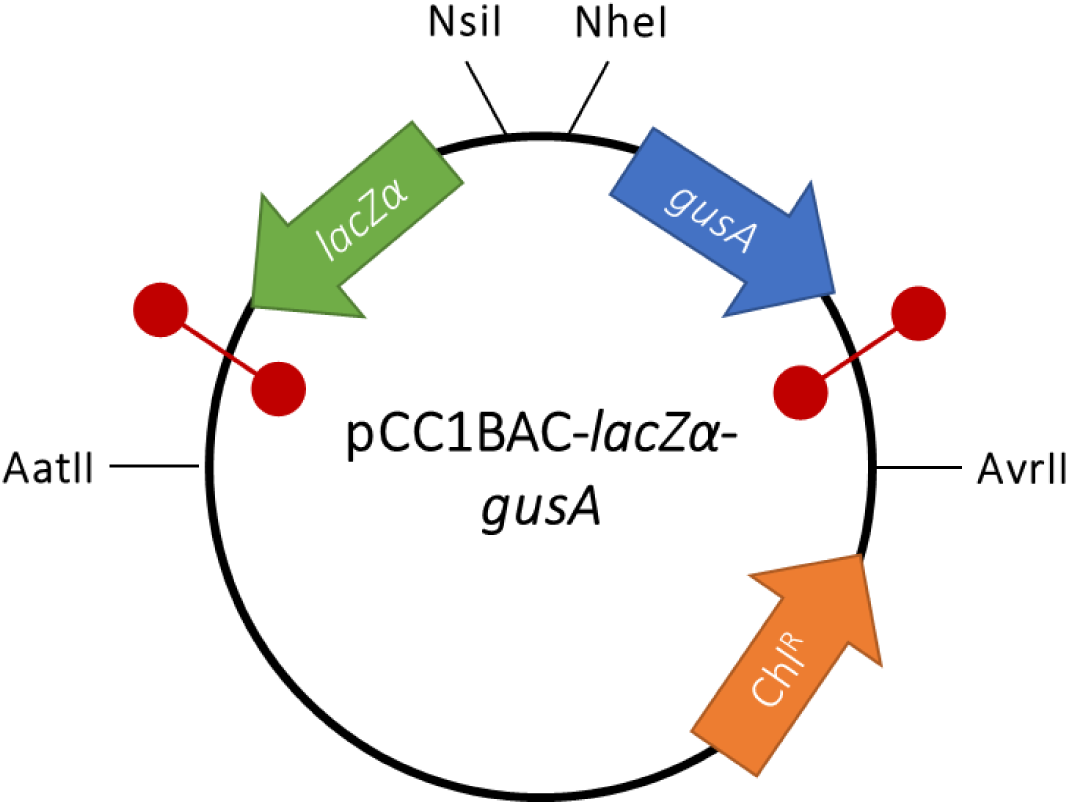
The structure of pCC1BAC-*lacZα*-*gusA*_plasmid. The green, blue and orange open arrowed boxes represent *lacZα, gusA* and chloramphenicol resistance gene, respectively, pointing in the direction of transcription. The black lines indicate the position of restriction sites on the plasmid. The red circles indicate bidirectional transcriptional terminators.

**Figure 3:**
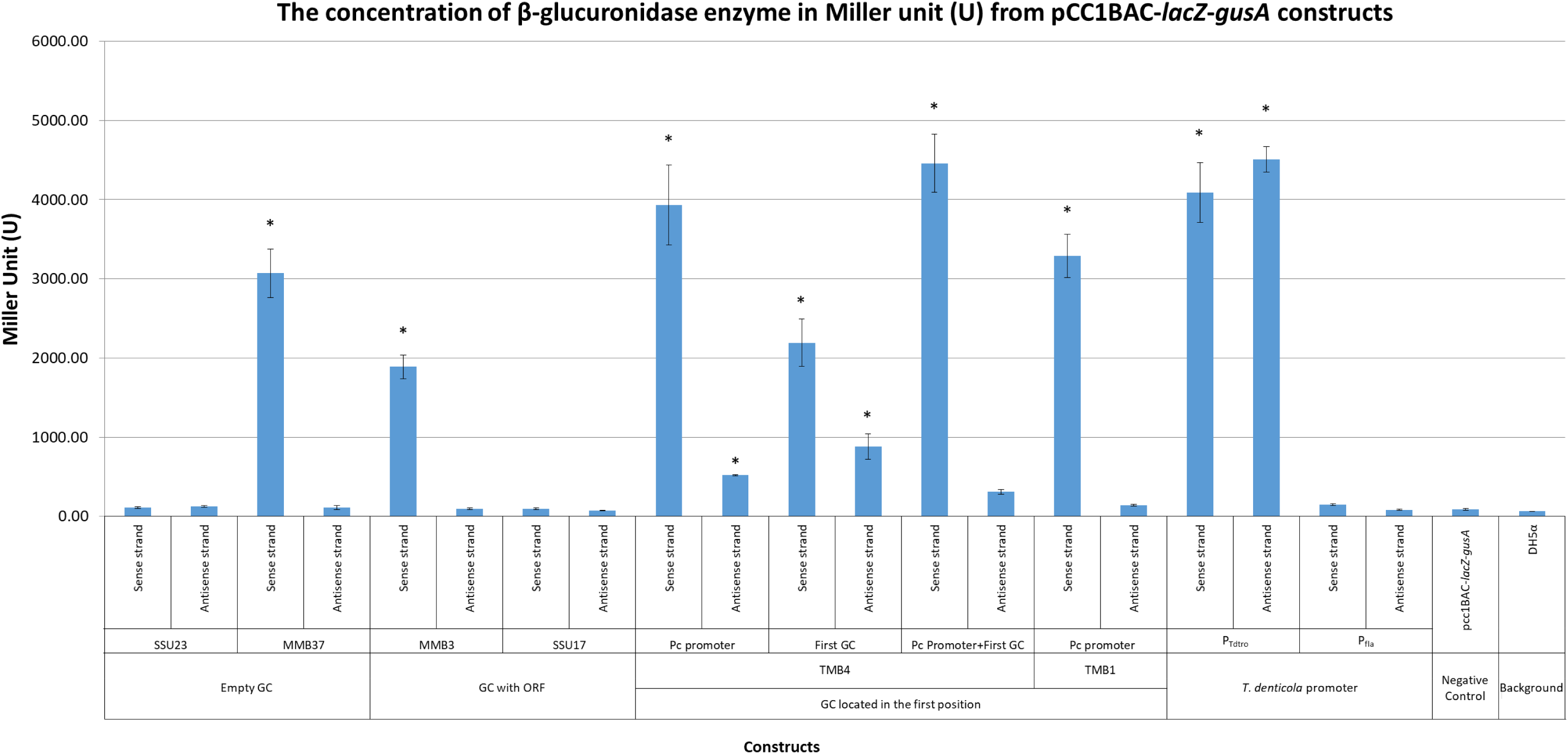
The promoter activity from pCC1BAC*-lacZα-GC-gusA* constructs estimated by β-glucuronidase enzyme assays. Error bars indicate the standard errors of the means from three replicates. The asterisks (*) indicate the constructs were statistically significantly different from the negative control group (pCC1BAC*-lacZα-gusA*) with the *p*-value <0.05 by using ordinary one-way ANOVA followed by Dunnett’s multiple comparison tests.

### Detection of promoter-containing GCs from oral metagenome

The p*CC1BAC-lacZα-gusA* plasmid, developed for the above enzymatic assay, had the potential to be used in an agar plate-based detection strategy to detect amplified integron GCs with promoter activity on either strand of DNA. This construct is called the Bi-Directional Promoter Detection plasmid (pBiDiPD). To verify the utility of pBiDiPD, and also to detect novel GCs containing promoter sequences in the human oral metagenome, integron GCs were amplified with *SUPA4-Nsi*I*/SUPA3-Nhe*I and *MARS5-Nsi*I*/MARS2-Nhe*I primers [28], and cloned into pBiDiPD. By spreading transformants on LB plates containing X-gal/IPTG and 4-methylumbelliferyl β-D-glucuronide (MUG), clones containing inserts with promoter activity in either direction could be identified. The clones with GCs containing a promoter on the sense strand showed blue fluorescence when visualised under UV light, reflecting the activity of β-glucuronidase enzymes catalysing MUG to yield the blue-fluorescent 4-methylumbelliferyl. Clones with promoter activity on the antisense strand resulted in blue colonies as a result of β-galactosidase enzymes catalysing X-Gal into a blue insoluble pigment 5,5’-dibromo-4,4’-dichloro-indigo (Fig 4).

**Figure 4:**
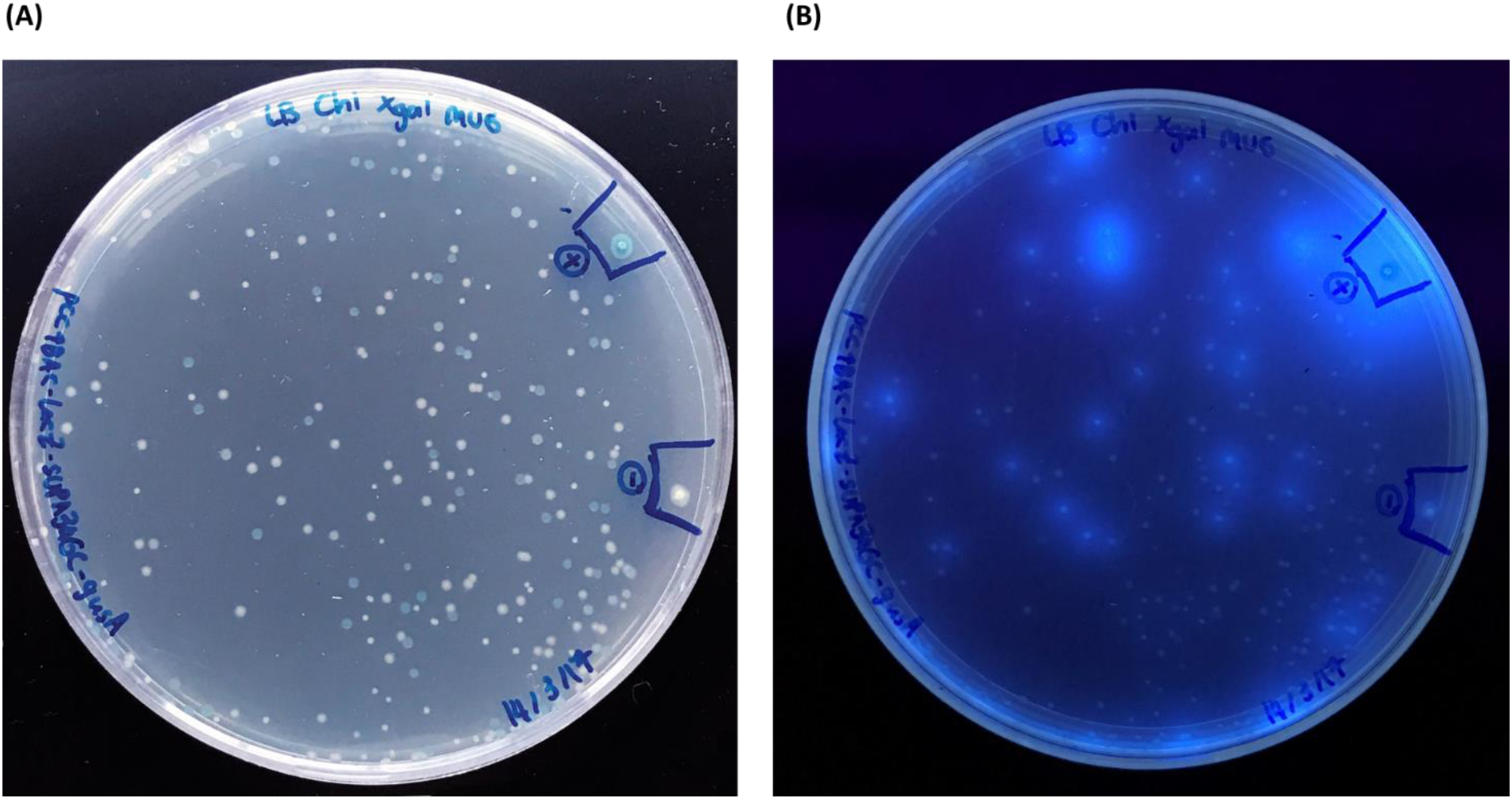
The detection of the integron GCs by using pBiDiPD. A.) Blue-white screening to detect for the clones with promoter activity on the antisense strand, B.) Exposing the colonies under the UV light to detect clones with promoter activity on the sense strand. The positive (+) and negative (−) colonies were the *E. coli* containing pCC1BAC-*lacZα*-TMB4-Pc-*gusA* (with experimentally proven promoter activities on either strand of DNA and pCC1BAC*-lacZα-gusA* (no promoter activity), respectively

After screening clones from both amplicon libraries (amplified with SUPA3-SUPA4 and MARS2-MARS5 primers), 23 different GCs with promoter activities were identified (Table 2). Fourteen of these were similar to the GCs identified in the previous study with >86% nucleotide identity [28]. Among the recovered promoter-containing GCs, 9 out of 23 were novel including sample SSU-Pro-20, SSU-Pro-27, SSU-Pro-32, SSU-Pro-46, SSU-Pro-65, MMU-Pro-5, MMU-Pro-24, and MMU-Pro-53. Artefactual PCRs were discounted by detecting the consensus R’ (1R) core sites [GTTRR(Y)R(Y)Y(R)] and the complementary R’’ (1L) core sites [R(Y)Y(R)Y(R)YAAC] of *attC* located downstream from the *attC* forward primers and upstream from the *attC* reverse primers, respectively (Supplementary Table 1) [35].

**Table 2.**
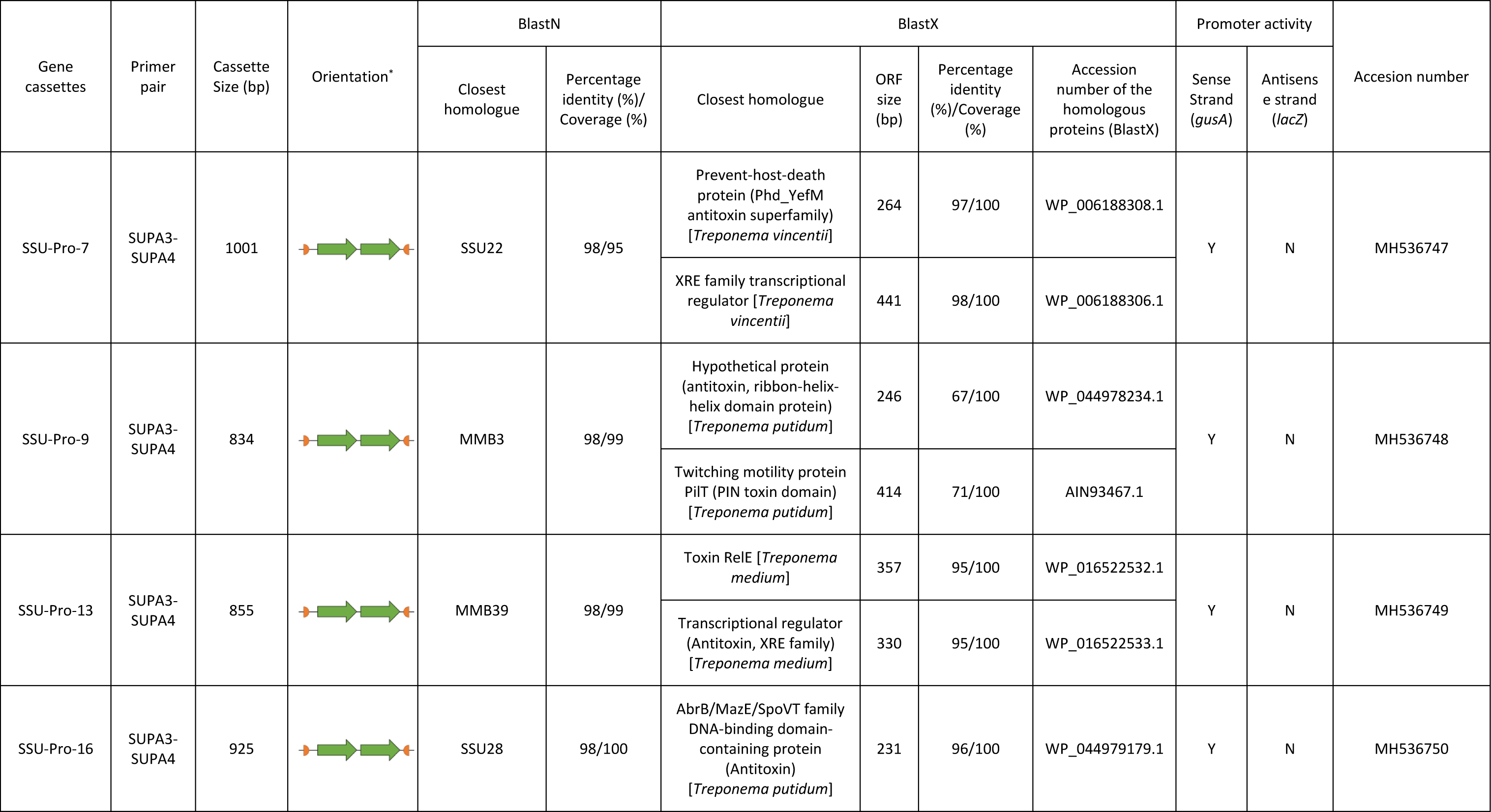
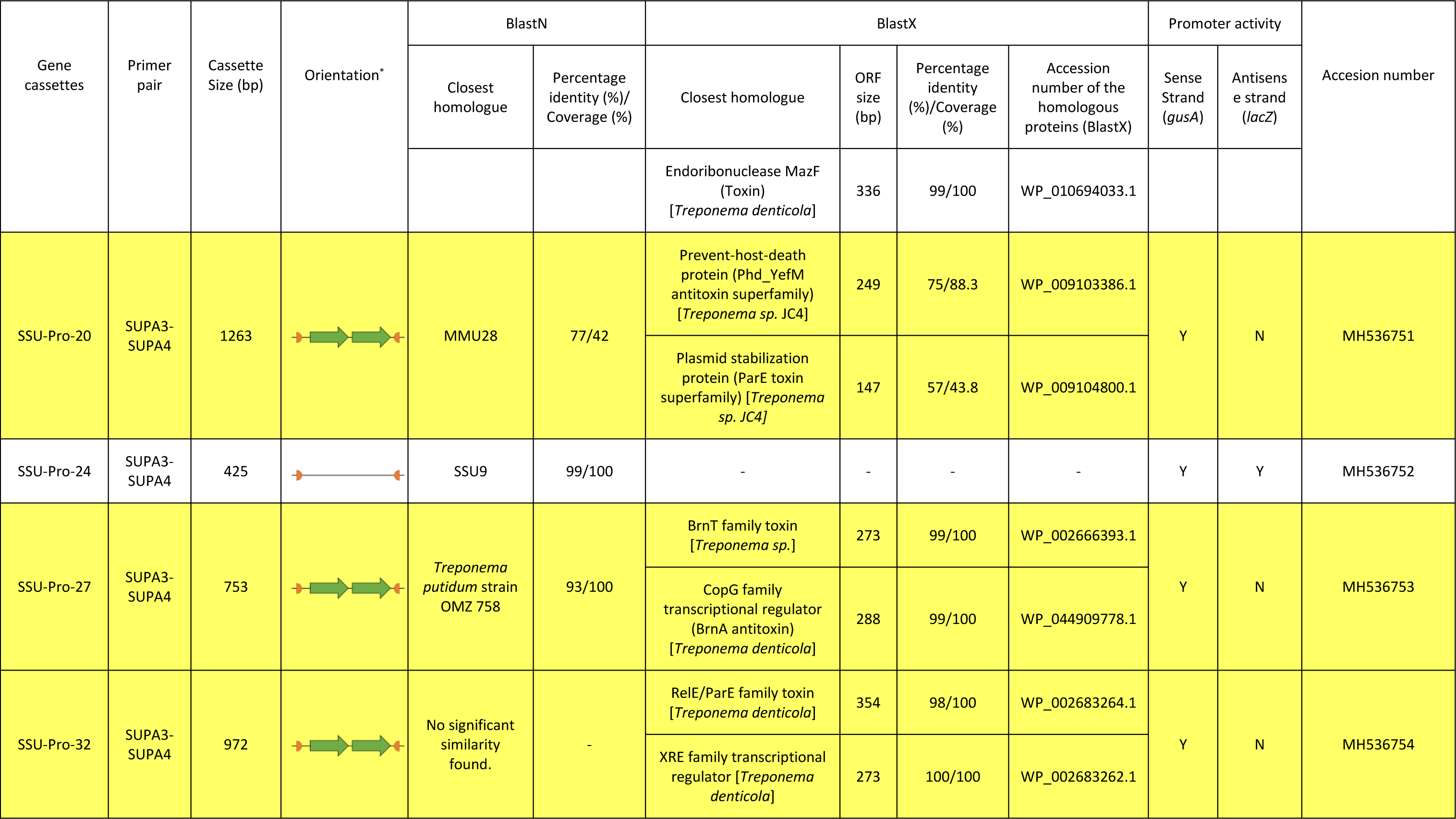
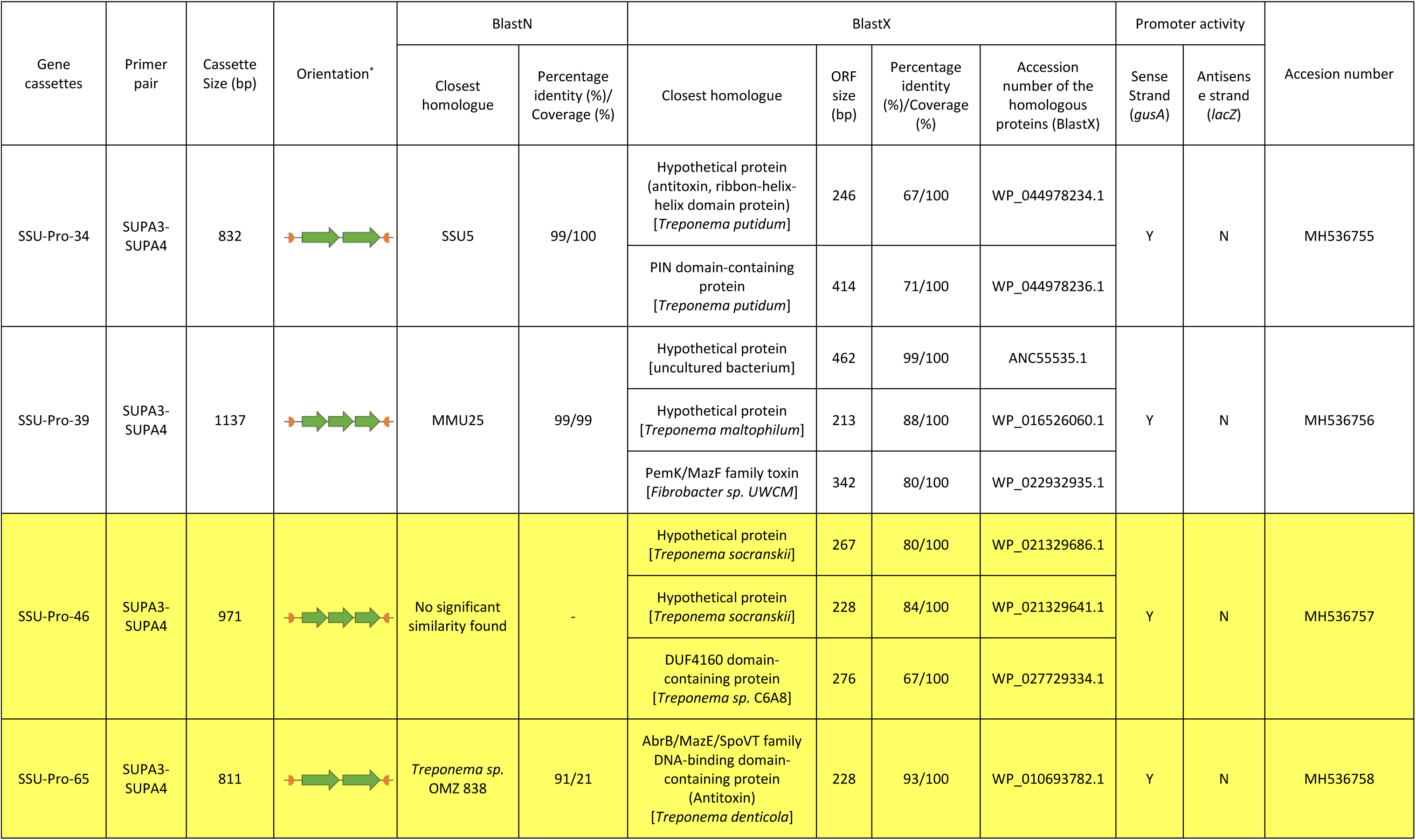
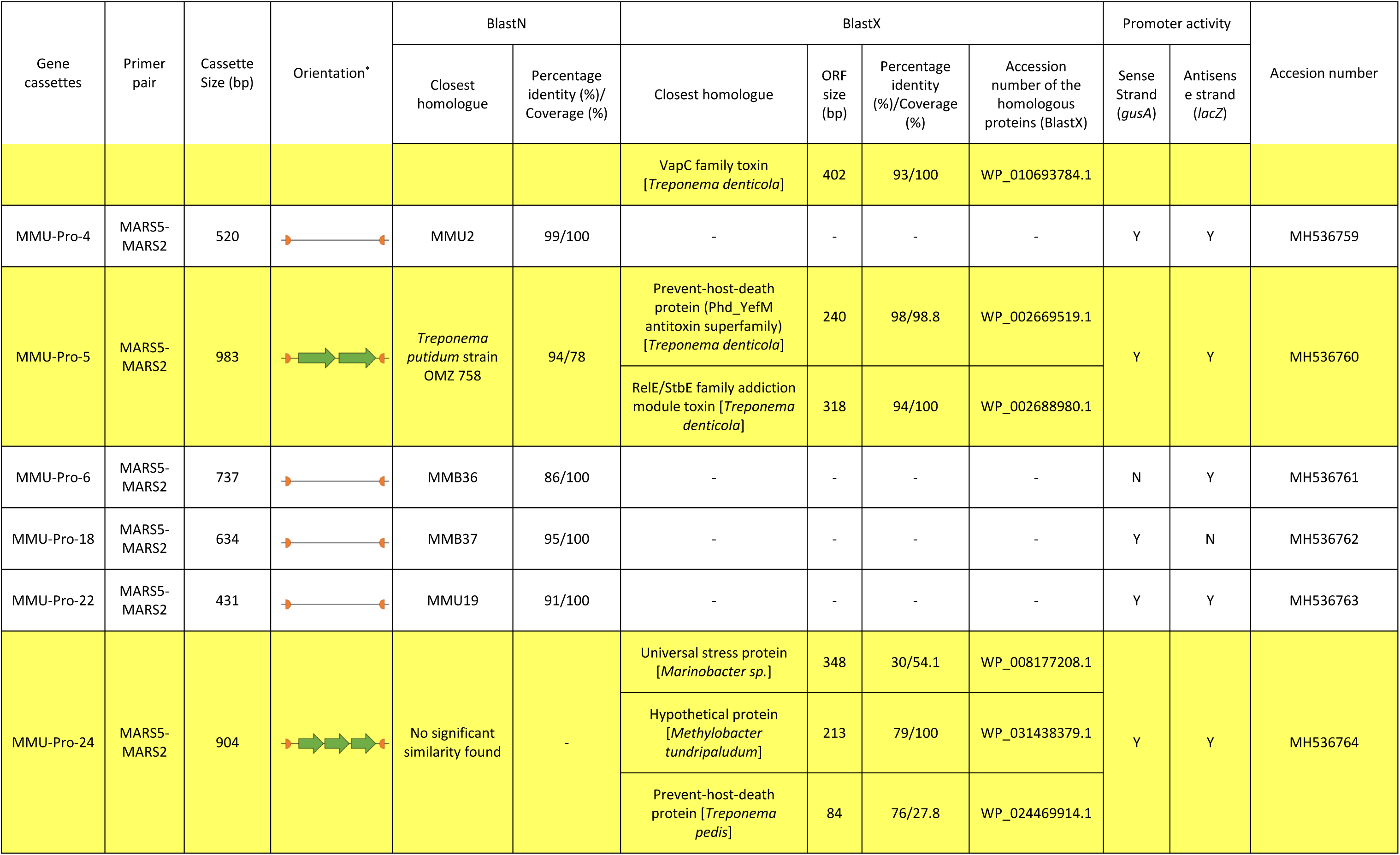
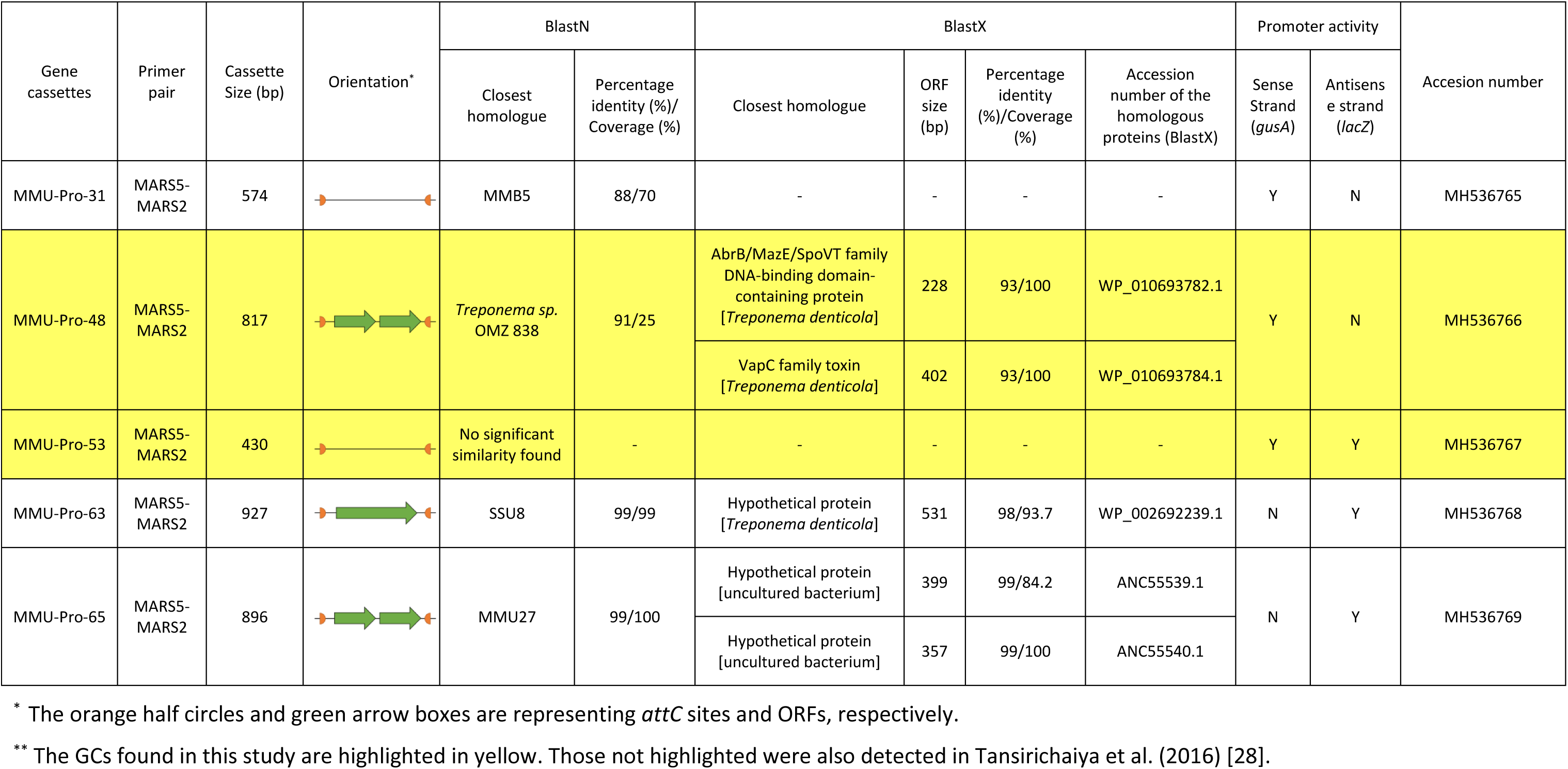
Characterisation of the human oral integron GCs containing promoter sequences detected by pBiDiPD.

| Gene cassettes | Primer pair | Cassette Size (bp) | Orientation* | BlastN |  | BlastX |  |  |  | Promoter activity |  | Accession number |
| --- | --- | --- | --- | --- | --- | --- | --- | --- | --- | --- | --- | --- |
|  |  |  |  | Closest homologue | Percentage identity (%)/ Coverage (%) | Closest homologue | ORF size (bp) | Percentage identity (%)/Coverage (%) | Accession number of the homologous proteins (BlastX) | Sense Strand (*gusA*) | Antisense strand (*lacZ*) |  |
| SSU-Pro-7 | SUPA3-SUPA4 | 1001 |  | SSU22 | 98/95 | Prevent-host-death protein (Phd_YefM antitoxin superfamily) [*Treponema vincentii*] | 264 | 97/100 | WP_006188308.1 | Y | N | MH536747 |
|  |  |  |  |  |  | XRE family transcriptional regulator [*Treponema vincentii*] | 441 | 98/100 | WP_006188306.1 |  |  |  |
| SSU-Pro-9 | SUPA3-SUPA4 | 834 |  | MMB3 | 98/99 | Hypothetical protein (antitoxin, ribbon-helix-helix domain protein) [*Treponema putidum*] | 246 | 67/100 | WP_044978234.1 | Y | N | MH536748 |
|  |  |  |  |  |  | Twitching motility protein PilT (PIN toxin domain) [*Treponema putidum*] | 414 | 71/100 | AIN93467.1 |  |  |  |
| SSU-Pro-13 | SUPA3-SUPA4 | 855 |  | MMB39 | 98/99 | Toxin RelE [*Treponema medium*] | 357 | 95/100 | WP_016522532.1 | Y | N | MH536749 |
|  |  |  |  |  |  | Transcriptional regulator (Antitoxin, XRE family) [*Treponema medium*] | 330 | 95/100 | WP_016522533.1 |  |  |  |
| SSU-Pro-16 | SUPA3-SUPA4 | 925 |  | SSU28 | 98/100 | AbrB/MazE/SpoVT family DNA-binding domain-containing protein (Antitoxin) [*Treponema putidum*] | 231 | 96/100 | WP_044979179.1 | Y | N | MH536750 |
|  |  |  |  | Closest homologue | Percentage identity (%)/Coverage (%) | Closest homologue | ORF size (bp) | Percentage identity (%)/Coverage (%) | Accession number of the homologous proteins (BlastX) | Sense Strand (*gusA*) | Antisense strand (*lacZ*) |  |
|  |  |  |  |  |  | Endoribonuclease MazF (Toxin) [*Treponema denticola*] | 336 | 99/100 | WP_010694033.1 |  |  |  |
| SSU-Pro-20 | SUPA3-SUPA4 | 1263 |  | MMU28 | 77/42 | Prevent-host-death protein (Phd_YefM antitoxin superfamily) [*Treponema sp.* JC4] | 249 | 75/88.3 | WP_009103386.1 | Y | N | MH536751 |
|  |  |  |  |  |  | Plasmid stabilization protein (ParE toxin superfamily) [*Treponema sp.* JC4] | 147 | 57/43.8 | WP_009104800.1 |  |  |  |
| SSU-Pro-24 | SUPA3-SUPA4 | 425 |  | SSU9 | 99/100 | - | - | - | - | Y | Y | MH536752 |
| SSU-Pro-27 | SUPA3-SUPA4 | 753 |  | *Treponema putidum* strain OMZ 758 | 93/100 | BrnT family toxin [*Treponema sp.*] | 273 | 99/100 | WP_002666393.1 | Y | N | MH536753 |
|  |  |  |  |  |  | CopG family transcriptional regulator (BrnA antitoxin) [*Treponema denticola*] | 288 | 99/100 | WP_044909778.1 |  |  |  |
| SSU-Pro-32 | SUPA3-SUPA4 | 972 |  | No significant similarity found. | - | RelE/ParE family toxin [*Treponema denticola*] | 354 | 98/100 | WP_002683264.1 | Y | N | MH536754 |
|  |  |  |  |  |  | XRE family transcriptional regulator [*Treponema denticola*] | 273 | 100/100 | WP_002683262.1 |  |  |  |
|  |  |  |  | Closest homologue | Percentage identity (%)/Coverage (%) | Closest homologue | ORF size (bp) | Percentage identity (%)/Coverage (%) | Accession number of the homologous proteins (BlastX) | Sense Strand (*gusA*) | Antisense strand (*lacZ*) |  |
| SSU-Pro-34 | SUPA3-SUPA4 | 832 |  | SSU5 | 99/100 | Hypothetical protein (antitoxin, ribbon-helix-helix domain protein) [*Treponema putidum*] | 246 | 67/100 | WP_044978234.1 | Y | N | MH536755 |
|  |  |  |  |  |  | PIN domain-containing protein [*Treponema putidum*] | 414 | 71/100 | WP_044978236.1 |  |  |  |
| SSU-Pro-39 | SUPA3-SUPA4 | 1137 |  | MMU25 | 99/99 | Hypothetical protein [uncultured bacterium] | 462 | 99/100 | ANC55535.1 | Y | N | MH536756 |
|  |  |  |  |  |  | Hypothetical protein [*Treponema maltophilum*] | 213 | 88/100 | WP_016526060.1 |  |  |  |
|  |  |  |  |  |  | PemK/MazF family toxin [*Fibrobacter sp. UWCM*] | 342 | 80/100 | WP_022932935.1 |  |  |  |
| SSU-Pro-46 | SUPA3-SUPA4 | 971 |  | No significant similarity found | - | Hypothetical protein [*Treponema socranskii*] | 267 | 80/100 | WP_021329686.1 | Y | N | MH536757 |
|  |  |  |  |  |  | Hypothetical protein [*Treponema socranskii*] | 228 | 84/100 | WP_021329641.1 |  |  |  |
|  |  |  |  |  |  | DUF4160 domain-containing protein [*Treponema sp. C6A8*] | 276 | 67/100 | WP_027729334.1 |  |  |  |
| SSU-Pro-65 | SUPA3-SUPA4 | 811 |  | *Treponema sp.* OMZ 838 | 91/21 | AbrB/MazE/SpoVT family DNA-binding domain-containing protein (Antitoxin) [*Treponema denticola*] | 228 | 93/100 | WP_010693782.1 | Y | N | MH536758 |
|  |  |  |  | Closest homologue | Percentage identity (%) / Coverage (%) | Closest homologue | ORF size (bp) | Percentage identity (%) / Coverage (%) | Accession number of the homologous proteins (BlastX) | Sense Strand (*gusA*) | Antisense strand (*lacZ*) |  |
|  |  |  |  |  |  | VapC family toxin [*Treponema denticola*] | 402 | 93/100 | WP_010693784.1 |  |  |  |
| MMU-Pro-4 | MARS5-MARS2 | 520 |  | MMU2 | 99/100 | - | - | - | - | Y | Y | MH536759 |
| MMU-Pro-5 | MARS5-MARS2 | 983 |  | *Treponema putidum* strain OMZ 758 | 94/78 | Prevent-host-death protein (Phd_YefM antitoxin superfamily) [*Treponema denticola*] | 240 | 98/98.8 | WP_002669519.1 | Y | Y | MH536760 |
|  |  |  |  |  |  | RelE/StbE family addiction module toxin [*Treponema denticola*] | 318 | 94/100 | WP_002688980.1 |  |  |  |
| MMU-Pro-6 | MARS5-MARS2 | 737 |  | MMB36 | 86/100 | - | - | - | - | N | Y | MH536761 |
| MMU-Pro-18 | MARS5-MARS2 | 634 |  | MMB37 | 95/100 | - | - | - | - | Y | N | MH536762 |
| MMU-Pro-22 | MARS5-MARS2 | 431 |  | MMU19 | 91/100 | - | - | - | - | Y | Y | MH536763 |
| MMU-Pro-24 | MARS5-MARS2 | 904 |  | No significant similarity found | - | Universal stress protein [*Marinobacter sp.*] | 348 | 30/54.1 | WP_008177208.1 | Y | Y | MH536764 |
|  |  |  |  |  |  | Hypothetical protein [*Methylobacter tundripaludum*] | 213 | 79/100 | WP_031438379.1 |  |  |  |
|  |  |  |  |  |  | Prevent-host-death protein [*Treponema pedis*] | 84 | 76/27.8 | WP_024469914.1 |  |  |  |
|  |  |  |  | Closest homologue | Percentage identity (%)/Coverage (%) | Closest homologue | ORF size (bp) | Percentage identity (%)/Coverage (%) | Accession number of the homologous proteins (BlastX) | Sense Strand (*gusA*) | Antisense strand (*lacZ*) |  |
| MMU-Pro-31 | MARS5-MARS2 | 574 |  | MMB5 | 88/70 | - | - | - | - | Y | N | MH536765 |
| MMU-Pro-48 | MARS5-MARS2 | 817 |  | *Treponema sp.* OMZ 838 | 91/25 | AbrB/MazE/SpoVT family DNA-binding domain-containing protein [*Treponema denticola*] | 228 | 93/100 | WP_010693782.1 | Y | N | MH536766 |
|  |  |  |  |  |  | VapC family toxin [*Treponema denticola*] | 402 | 93/100 | WP_010693784.1 |  |  |  |
| MMU-Pro-53 | MARS5-MARS2 | 430 |  | No significant similarity found | - | - | - | - | - | Y | Y | MH536767 |
| MMU-Pro-63 | MARS5-MARS2 | 927 |  | SSU8 | 99/99 | Hypothetical protein [*Treponema denticola*] | 531 | 98/93.7 | WP_002692239.1 | N | Y | MH536768 |
| MMU-Pro-65 | MARS5-MARS2 | 896 |  | MMU27 | 99/100 | Hypothetical protein [uncultured bacterium] | 399 | 99/84.2 | ANC55539.1 | N | Y | MH536769 |
|  |  |  |  |  |  | Hypothetical protein [uncultured bacterium] | 357 | 99/100 | ANC55540.1 |  |  |  |

\* The orange half circles and green arrow boxes are representing *attC* sites and ORFs, respectively.
\*\* The GCs found in this study are highlighted in yellow. Those not highlighted were also detected in Tansirichaiya et al. (2016) [28].

The GCs can be categorised into two groups, one predicted to encode toxin-antitoxin systems in 12 out of 23 GCs, including plasmid stabilization protein (toxin)-prevent-host-death protein (antitoxin), BrnT (toxin)-BrnA (antitoxin), VapC (toxin)-AbrB/MazE/SpoVT family protein (antitoxin), RelE/ParE family (toxin)-XRE transcriptional regulator (antitoxin). The second group contained ORF-less GCs, which could be found in 7 samples, all reported in the previous study, except sample MMU-Pro-53. Most of the samples (14 out of 23 GCs) showed the promoter activity only on the sense strand. Samples with promoter activity only on the antisense strand were MMU-Pro-6, MMU-Pro-63, and MMU-Pro-65, while 6 out of 23 GCs exhibited promoter activity on both strands.

## Discussion

Integrons are important disseminators of antimicrobial resistance genes and therefore, it is important to understand the diversity of GCs and how their expression is controlled. Even though most of the GCs contained a single ORF, ORF-less GCs have also been found [24, 27, 28, 36, 37].

In this study, we determined promoter activity from GCs isolated by PCR from metagenomic DNA by measuring promoter activity from multiple GC containing constructs. As the ORF-less GCs were recovered from the oral metagenome, there is little information regarding the original host. Therefore, we chose to test the promoter activities by using an *E. coli* surrogate. Nucleotide sequence analysis suggested that these GCs were likely to be derived from *Treponema* spp., therefore, the ability of *E. coli* to utilise *T. denticola* promoter sequences was determined by including the experimentally verified *T. denticola* promoter, PTdTro [31] which showed high activity on both sense and antisense strands, providing confidence that *E. coli* could be used. However, as no promoter activity was detected from P_Fla_, it suggested that our enzymatic assay cannot detect promoters associated with σ^28^ from *Treponema* spp., which could be due to an inability for the *E. coli* host to recognise the *Treponema* σ^28^ promoter sequence.

Promoter activities of the ORF-less GCs were confirmed and quantified by using a β-glucuronidase enzymatic assay. This is the first time that the promoter activity of ORF-less GCs has been demonstrated *in vitro*, as shown by the activity on the sense strand of the MMB37 and both strands of the TMB4. A study on the *Vibrio* integron, containing a 116-cassette array, showed that most of the GCs are transcribed [38]. Therefore, ORF-less GCs could be responsible for transcription of the other GCs not transcribed by P_C_.

For the TMB4 GC (ORF-less GC in the first position), it was initially hypothesised that the promoter could help increase the expression of the downstream GCs. This was shown when P_C_ promoter was coupled with a second promoter (P_2_) (found in 10% of class 1 integron and located 119 bp downstream from P_C_), could result in a significantly higher expression of GCs [39, 40]. The constructs of TMB4 P_C_ and TMB4 P_C_+GC were therefore included in the assay to determine whether having a promoter GC at the first position could help promote the expression of downstream GCs. The results show that coupling promoter GC in the first position slightly increased the expression of reporter genes (Fig. 3). However, this was not significant (*p*-value >0.99 by using ordinary one-way ANOVA followed by Bonferroni’s post-hoc).

The lack of additive promoter activity can be explained by more competition for enzymes involved in transcription such as RNA polymerases (RNAP) or sigma factors to be available for transcription from each promoter, resulting in lower transcriptional level [41]. Another, not mutually exclusive possibility is transcriptional interference (TI) between the four promoters on the TMB4 P_C_+GC construct. We have experimentally shown promoter activity of TMB4 P_C_ and TMB4 GC constructs on both strands, indicating convergent TI is a possibility.

In usual integrons, P_C_ is in *intI*, which is convergent to the integron integrase promoter P*_IntI_* downstream (Fig 1), resulting in TI. The TI between P_C_ and P*_IntI_* has been shown to control the expression of integrase and the subsequent recombination of GCs. The weaker strength of P_C_ could result in higher expression of integrase, which increases recombination of GCs [42, 43]. This relationship of P_C_ and *intI* might also apply to the reverse integrons found in *T. denticola*, even though their position and direction of *P_intI_*, P_C_ and *intI* gene are in a different orientation compared to the usual integrons (Fig 1).

Due to the lack of additive promoter activity when Pc and an ORF-less GC promoter were tested in tandem we hypothesised that there is an alternative selective advantage for having an ORF-less, promoter-containing GC in the first position on an integron GC array.

The expression level of cassette genes located further down in the array normally decreases due to the formation of a stem-loop structure on mRNA at *attC* sites, which impede the progression of the ribosome [44]. It was previously shown that the level of streptomycin resistance was reduced four times, when the *aadA2*-containing GC was located in the second position [45]. However, our data shows that the insertion of an ORF-less, promoter-containing GC in the first position did not decrease the *gusA* expression significantly (considered as a proxy for the expression of gene(s) in the second GC), i.e. comparing the data for TMB4 P_C_ and TMB4 P_C_+GC. Therefore, we hypothesised that promoter-containing GCs could act as a genetic clutch, where the expression of the original first GC is partially disengaged from the P_C_ promoter and replaced by the one on the ORF-less promoter containing GC (Fig 5A). This can prevent a significant change in expression of the first GC while a new, first GC is sampled from the pool of GCs in order to adapt to an additional stress concurrent with the selective pressure requiring expression of the first GC. This system would work as a genetic clutch with the insertion of any GC containing a promoter in the same direction as P_C_, so it could be the insertion of either ORF-less GCs such as TMB4 GC, or other promoter-containing GCs such as the multiple TA-containing GCs we have identified; providing another selective advantage to retaining them and explaining their varied position within the GC array.

**Figure 5:**
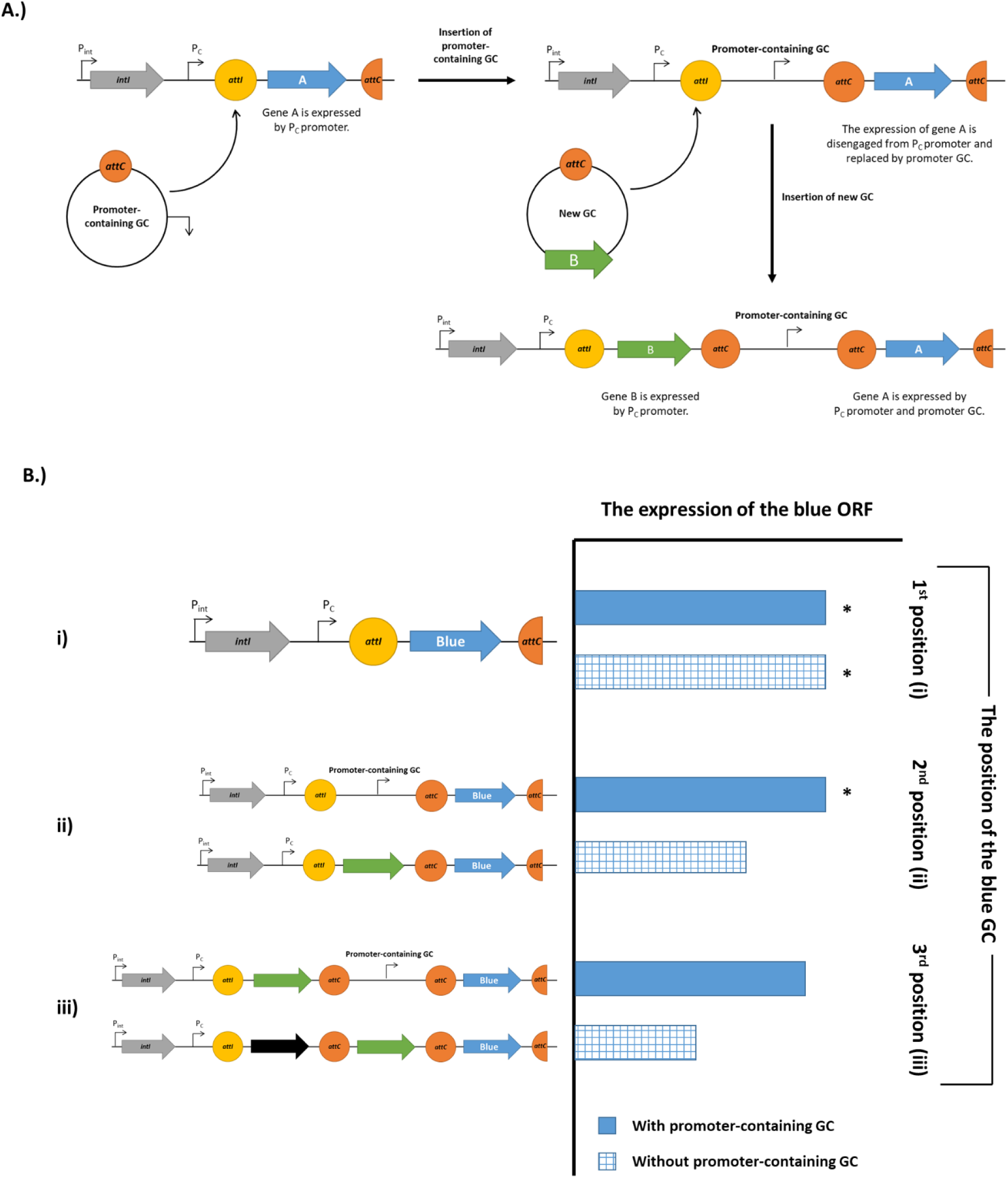
The proposed genetic clutch. (A) When a promoter-containing GC inserts into the first position, it can act as a genetic clutch by disengaging the original first GC (blue arrow) from P_C_ promoter and replaced with the one on promoter GC. When a new GC (green arrow) inserts, it can be expressed by P_C_ promoter, while the blue GC is expressed by promoter-containing GC and P_C_ promoter. (B) The expression level of gene cassettes with and without a genetic clutch. The estimated levels of expression of the blue ORF in i.) the first, ii.) the second and iii.) the third position were shown in the bar chart. The solid bars represent the situation when promoter-containing GC was inserted upstream of the blue GC, while the gridded bars represent the situation when no promoter-containing GC was inserted. The asterisks indicate the experimentally verified expression level, suggested by the results in Figure 3 (TMB4 P_C_ and TMB4 P_C_+GC). The expression of the blue ORF was hypothesised to be decreased when more GCs are inserted without the presence of a promoter-containing GC as a genetic clutch (gridded bars), based on the data from previous study [45].

A genetic clutch within an integron can be of benefit to bacteria when they are exposed to multiple environmental stresses such as two different antibiotics simultaneously. The first resistance gene (green ORF in Fig 5Biii) can be expressed by the P_C_ promoter, while the second resistance gene (blue GC), located in the third position, is expressed by P_C_ and the promoter GC. Therefore, allowing bacteria to survive in both the presence of both drugs.

As the other ORF-less GC MMU23 showed no promoter activity it may have other functions or carry a promoter that can be recognised in its native host but not in *E. coli*, or require other sigma factors. For the ORF-containing GC MMB3 sample, the promoter activity was found on the sense strand. This GC was predicted to carry toxin-antitoxin (TA) ORFs, including the PIN toxin and ribbon-helix-helix antitoxin domains, which were shown to contain their own promoter. Sample SSU17 and MMU23, which showed no promoter activity, can be considered as a control; illustrating that not all of GCs amplified from the oral metagenome exhibited promoter activity within our assay.

The p*CC1BAC-lacZα-gusA* plasmid, developed for the enzymatic assay, also had potential to be used for the detection of promoter activity in either direction from GCs. The clones with promoters on the sense strand can be detected under UV light and showed blue fluorescence because β-glucuronidase can cleave the substrate, MUG, on the plate, which produces a fluorescence compound called methylumbelliferone. For the clones carrying promoters on the antisense strand, they can be detected by blue-white screening as β-galactosidase can cleave X-gal, producing an intensively blue product called 5,5’-dibromo-4,4’-dichloro-indigo, which can be viewed by eye under normal light.

To verify the application of pCC1BAC-*lacZα-gusA* plasmids as promoter detection system, integron GCs were amplified from the human saliva metagenome by using SUPA3-SUPA4 and MARS2-MARS5 primers, which amplify integron GCs from the oral metagenome [28]. After cloning the amplified GCs between both reporter genes, two groups of GCs were identified with promoter activities: ORF-less GCs and TA-containing GCs. By detecting 7 clones containing ORF-less GCs with promoter activity it further supported that one of the functions of ORF-less GCs in integrons is to provide promoter activities.

TA-containing GCs are abundant in chromosomal integrons (CIs), which were suggested to have a role in preventing random deletion of GCs and stabilising the large arrays CIs [22, 46, 47]. TA systems normally encode a stable toxin and a labile antitoxin [48], therefore TA cassettes have to carry their own promoters to ensure their expression. These were found in CIs of *Treponema* spp., such as the HicA-HicB TA-containing GC in the fourth position within the GC array (Accession number NC_002967) in the CI from *T. denticola* [30]. As most of the GCs amplified with our primers were homologous with *Treponema* spp., these TA-containing GCs should be present in our oral metagenome and were detected by our pBiDiPD based on their promoter activities.

Two of the GCs, SSU-Pro-9 and MMU-Pro-18, were similar to the MMB3 and MMB37 GCs, respectively, which were shown by the β-glucuronidase enzyme assay to have promoter activity on the sense strand. The phenotypes of SSU-Pro-9 and MMU-Pro-18 colonies also showed only a blue fluorescence phenotype, reflecting the promoter activity on the sense strand, which corresponded with the enzymatic assay results of MMB3 and MMB37.

To summarise, the promoter activities of the ORF-less integron GCs were experimentally demonstrated by using a robust β-glucuronidase enzyme assay, confirming that one of the functions of ORF-less GCs is to provide promoters for the expression of ORF containing GCs, in addition to expression from P_C_. The dual reporter plasmid; pBiDiPD, was developed for the direct visualisation of clones containing gene cassettes with promoter activity on agar plates. This can be applied as a detection system for promoter activity for any other DNA fragments.

## Materials and methods

### *in silico* analysis of the human oral cavity gene cassettes and the construction of pCC1BAC-*lacZα*-GC-*gusA* constructs

All of the ORF-less GCs and some of the GCs containing ORFs, identified in the previous study [28], were predicted for putative promoter sequences by using the web-based software BPROM in the Softberry package [49].

### Construction of pUC19-GC-*gusA* and pCC1BAC*-lacZα-GC-gusA* constructs

To determine the promoter activity of the selected GCs, the constructs were initially cloned into the EcoRI and KpnI restriction sites on pUC19-P*tet*(M)-*gus*A plasmid [50]. The selected GCs were amplified from the pGEM-T easy vector containing the GC amplicon from a previous study [28], as shown in Supplementary Fig 1, by using primer listed in Supplementary Table 2.

Due to a significant difference in the plasmid copy number in some constructs of the pUC19-GC-*gusA*, new constructs were prepared based on a low copy number CopyControl™ pCC1BAC™ vector (Epicenter, UK) as it will be maintained in *E. coli* cell as one plasmid per cell and enable us to control the plasmid copy number to be similar between each construct. The construct was designed to contain two reporter genes, β-galactosidase *lacZα* and β-glucuronidase *gusA* genes (Fig 2 and Supplementary Fig 2). As *lacZα* on pCC1BAC contained T7 promoter sequence, it was first deleted by using Q5^®^ Site-Directed Mutagenesis Kit (New England Biolabs, UK). The backbone of pCC1BAC was amplified with pCC1BAC-del*LocZ*-F1 and pCC1BAC-del*LocZ*-R1, and the amplified products were treated with a Kinase-Ligase-DpnI (KLD) enzyme mix, following the instructions from the manufacturer. The KLD-treated product was then transformed into *E. coli* α-Select Silver Efficiency competent cells (Bioline, UK) following the instructions from the manufacturer. The pCC1BAC-del*LacZ* plasmid was then extracted from *E. coli* by using QIAprep Spin Miniprep Kit (Qiagen, UK), following the manufacturer’s instructions.

The *lacZα* reporter gene was amplified from the pUC19 vector (New England Biolabs, UK) with *LacZ-*F1 and *LacZ-*R1 primers. For *gusA* reporter gene, it was amplified from pUC19-*Ptet*(M)-*gusA* with *gusA-F1* and *gusA-R1* primers. A bidirectional terminator, modified from *lux* operon, was added to *LacZ-F1* and *gusA*-R1 primers, resulting in two bi-directional terminators flanking the *lacZα-gusA* reporter genes [51]. This was done to prevent transcriptional read-through from the promoter in the plasmid backbone and to also prevent promoters from the inserts interfering with the expression of genes on the plasmid backbone. The *lacZα* and *gusA* amplicons were digested with NsiI restriction enzymes (New England Biolabs, UK) and ligated together by using T4 DNA ligase (New England Biolabs, UK). The *lacZα-gusA* ligated product was directionally cloned into the pCC1BAC-delLacZ plasmid by digesting them with AatII and AvrII restriction enzymes and ligated together, resulting in pCC1BAC*-lacZα-gusA* plasmid.

The selected GCs were amplified from each pUC19-GC-*gusA* constructs by using primer listed in Supplementary Table 1. The amplicons were double digested with *Nsi*I and *Nhe*I and directionally cloned into a pre-digested pCC1BAC*-lacZα-gusA* plasmid, then transformed into *E. coli* α-Select Silver Efficiency competent cells.

### Determination of β-glucuronidase enzymatic activity

The β-glucuronidase enzymatic assay was performed to measure the promoter activity based on the expression of *gusA*, following the protocol described previously with some modifications [52]. The overnight cultures of *E. coli* containing the reporter constructs were prepared in LB broth supplemented with 12.5 μg/mL chloramphenicol. The OD_600_ of each overnight culture was measured. An aliquot of 1 mL of the overnight culture was centrifuged at 3000 × *g* for 10 min and discarded the supernatant. The cell pellets were incubated at −70°C for 1 hr and resuspended in 800 μl of pH 7 Z buffer (50 mM 2-mercaptoethanol, 40 mM NaH_2_PO_4_·H_2_O, 60 mM Na_2_HPO_4_·7H_2_O, 10 mM KCl, and 1mM MgSO_4_·7H_2_O) and 8 μl of toluene. The mixture was transferred to a 2 ml cryotube containing glass beads (150–212 μm in diameter) (Sigma, UK) and vortexed twice for 5 min each with an incubation on ice for 1 min in between. The glass beads were then removed by centrifugation at 3000 × *g* for 3 min. One-hundred microliters of cell lysate were mixed with 700 μl of Z-buffer, then incubated at 37°C for 5min. One-hundred sixty microliters of 6 mM ρ-nitrophenyl-β-D-glucuronide (PNPG) was then added to the reaction and incubated at 37°C for 5 min. The reactions were stopped by adding 400 μl of 1 M Na_2_CO_3_ and centrifuged at 3000 × *g* for 10 min to remove cell debris and glass beads. The absorbance of the supernatant was measured with a spectrophotometer at the wavelength of 405 nm. Three biological replicates of the β-glucuronidase enzymatic assay were performed. The β-glucuronidase Miller units were calculated from 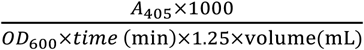 [53].

### Statistical analysis

The average and standard deviation of β-glucuronidase concentration were calculated from three biological replicates, which were used for the columns and error bars in figure 3, respectively. The statistical comparisons between the negative control (pCC1BAC*-lacZ-gusA*) to the other constructs were performed by using ordinary one-way ANOVA with either Dunnett’s post-hoc test (to compare each construct with a negative control) or Bonferroni’s post-hoc test (to compare constructs between themselves). The groups with statistically significantly difference from the control had the *p*-value of less than 0.05.

### Recovery of promoter-containing GCs from the human oral metagenome

The integron GCs were amplified from the human oral metagenome by using as described previously with SUPA4-NsiI-SUPA4-NheI and MARS5-NsiI-MARS2-NheI primers [28]. The human oral metagenomic DNA was previously extracted from the saliva samples collected from 11 volunteers in the Department of Microbial Diseases, UCL Eastman Dental Institute [28]. The Ethical approval for the collection and uses of saliva samples was obtained from University College London (UCL) Ethics Committee (project number 5017/001).

The amplified products were purified and digested with NsiI and NheI and ligated with the pre-digested pCC1BAC*-lacZα-gusA* plasmid. The ligated products were transformed into *E. coli* α-Select Silver Efficiency competent cells by heat shock. Cells were spread on LB agar supplemented with 12.5 μg/mL chloramphenicol, 80 μg/mL X-gal, 50μM IPTG, and 70 μg/mL 4-methylumbelliferyl-β-D-glucuronide (MUG). After incubation at 37°C for 18 hr, the colonies with β-galactosidase activity from *lacZ* was detected by blue-white screening on the agar plate, and the β-glucuronidase activity from *gusA* was visualisation under UV light. Colonies exhibiting either activity were selected and subcultured on fresh agar plates. The inserts were amplified by colony PCR using *lacZ-*F2 and *gusA-*F2 primers and sequenced by sequencing service from Genewiz (Genewiz, UK).

### Sequence analysis and nomenclature of promoter-containing GC amplicons

DNA sequences were visualised and analysed by using BioEdit version 7.2.0 (http://www.mbio.ncsu.edu/bioedit/bioedit.html). The contigs from sequencing reactions were combined by using CAP contig function in the software. The sequences were then matched to the nucleotide and protein database by using BlastN and BlastX from the National Centre for Biotechnology Information (NCBI), respectively. The criteria for the sequence analysis of integron GC were the same as described in the previous study [28]. Two additional criteria for the verification of GCs detected with pCC1BAC*-lacZα-gusA* were included. Any clones containing incomplete GCs, caused by digestion at internal NsiI and NheI restriction sites on the GCs, were excluded from the dataset. Also chimeric inserts, which were the ligation products between digested amplicons, were also excluded.

The promoter-containing GCs were named as described in the previous study [28]. The first and second letters represented the forward primer and reverse primer used in the amplification. The third letter represents the source of the human oral metagenomic DNA which is U for the United Kingdom. This was followed by term “Pro”, indicating the presence of promoter activity, and the number of the clone. For instance, SSU-Pro-1 stands for the first clone amplified from the UK oral metagenome by using SUPA3 and SUPA4 primers. The sequences of these GCs were deposited in the DNA database with the accession number from MH536747 to MH536769.

## References

1. Hall RM, Collis CM. Antibiotic resistance in gram-negative bacteria: the role of gene cassettes and integrons. Drug resistance updates : reviews and commentaries in antimicrobial and anticancer chemotherapy. 1998;1(2):109–19. Epub 2006/08/15. PubMed PMID: 16904397.

2. Cambray G, Guerout AM, Mazel D. Integrons. Annual review of genetics. 2010;44:141–66. Epub 2010/08/17. doi: 10.1146/annurev-genet-102209-163504. PubMed PMID: 20707672.

3. Michael CA, Gillings MR, Holmes AJ, Hughes L, Andrew NR, Holley MP, et al. Mobile gene cassettes: a fundamental resource for bacterial evolution. The American naturalist. 2004;164(1):1–12. Epub 2004/07/22. doi: 10.1086/421733. PubMed PMID: 15266366.

4. Escudero JA, Loot* C, Nivina A, Mazel D. The Integron: Adaptation On Demand. Microbiology spectrum. 2015;3(2). doi: doi:10.1128/microbiolspec.MDNA3-0019-2014.

5. Boucher Y, Labbate M, Koenig JE, Stokes HW. Integrons: mobilizable platforms that promote genetic diversity in bacteria. Trends in microbiology. 2007;15. doi: 10.1016/j.tim.2007.05.004.

6. Wu Y-W, Doak TG, Ye Y. The gain and loss of chromosomal integron systems in the *Treponema* species. BMC evolutionary biology. 2013;13(1):1–9. doi: 10.1186/1471-2148-13-16.

7. Mazel D. Integrons: agents of bacterial evolution. Nature reviews. 2006;4.

8. Gillings MR. Integrons: past, present, and future. Microbiology and molecular biology reviews : MMBR. 2014;78(2):257–77. Epub 2014/05/23. doi: 10.1128/mmbr.00056-13. PubMed PMID: 24847022; PubMed Central PMCID: PMCPMC4054258.

9. Collis CM, Hall RM. Gene cassettes from the insert region of integrons are excised as covalently closed circles. Molecular microbiology. 1992;6(19):2875–85. Epub 1992/10/01. PubMed PMID: 1331702.

10. Partridge SR, Recchia GD, Scaramuzzi C, Collis CM, Stokes HW, Hall RM. Definition of the *attI1* site of class 1 integrons. Microbiology (Reading, England). 2000;146 ( Pt 11):2855–64. Epub 2000/11/07. PubMed PMID: 11065364.

11. Collis CM, Recchia GD, Kim M-J, Stokes HW, Hall RM. Efficiency of Recombination Reactions Catalyzed by Class 1 Integron Integrase IntI1. Journal of bacteriology. 2001;183(8):2535–42. doi: 10.1128/JB.183.8.2535-2542.2001. PubMed PMID: PMC95170.

12. Guerin E, Cambray G, Sanchez-Alberola N, Campoy S, Erill I, Da Re S, et al. The SOS response controls integron recombination. Science (New York, NY). 2009;324(5930):1034. Epub 2009/05/23. doi: 10.1126/science.1172914. PubMed PMID: 19460999.

13. Baharoglu Z, Bikard D, Mazel D. Conjugative DNA transfer induces the bacterial SOS response and promotes antibiotic resistance development through integron activation. PLOS Genetics. 2010;6(10):e1001165. doi: 10.1371/journal.pgen.1001165.

14. Baharoglu Z, Krin E, Mazel D. Connecting environment and genome plasticity in the characterization of transformation-induced SOS regulation and carbon catabolite control of the *Vibrio cholerae* integron integrase. Journal of bacteriology. 2012;194(7):1659–67. Epub 2012/01/31. doi: 10.1128/jb.05982-11. PubMed PMID: 22287520; PubMed Central PMCID: PMCPMC3302476.

15. Baharoglu Z, Mazel D. *Vibrio cholerae* triggers SOS and mutagenesis in response to a wide range of antibiotics: a route towards multiresistance. Antimicrobial agents and chemotherapy. 2011;55(5):2438–41. Epub 2011/02/09. doi: 10.1128/aac.01549-10. PubMed PMID: 21300836; PubMed Central PMCID: PMCPMC3088271.

16. Little JW. Mechanism of specific LexA cleavage: autodigestion and the role of RecA coprotease. Biochimie. 1991;73(4):411–21. Epub 1991/04/01. PubMed PMID: 1911941.

17. Harms K, Starikova I, Johnsen PJ. Costly Class-1 integrons and the domestication of the the functional integrase. Mobile genetic elements. 2013;3(2):e24774. Epub 2013/08/06. doi: 10.4161/mge.24774. PubMed PMID: 23914313; PubMed Central PMCID: PMCPMC3681742.

18. Starikova I, Harms K, Haugen P, Lunde TT, Primicerio R, Samuelsen O, et al. A trade-off between the fitness cost of functional integrases and long-term stability of integrons. PLoS Pathog. 2012;8(11):e1003043. Epub 2012/12/05. doi: 10.1371/journal.ppat.1003043. PubMed PMID: 23209414; PubMed Central PMCID: PMCPMC3510236.

19. Coyne S, Guigon G, Courvalin P, Perichon B. Screening and quantification of the expression of antibiotic resistance genes in *Acinetobacter baumannii* with a microarray. Antimicrobial agents and chemotherapy. 2010;54(1):333–40. Epub 2009/11/04. doi: 10.1128/aac.01037-09. PubMed PMID: 19884373; PubMed Central PMCID: PMCPMC2798560.

20. Stokes HW, Hall RM. Sequence analysis of the inducible chloramphenicol resistance determinant in the *Tn1696* integron suggests regulation by translational attenuation. Plasmid. 1991;26(1):10–9. Epub 1991/07/01. PubMed PMID: 1658833.

21. da Fonseca EL, Vicente AC. Functional characterization of a Cassette-specific promoter in the class 1 integron-associated *qnrVC1* gene. Antimicrobial agents and chemotherapy. 2012;56(6):3392–4. Epub 2012/03/07. doi: 10.1128/aac.00113-12. PubMed PMID: 22391535; PubMed Central PMCID: PMCPMC3370728.

22. Szekeres S, Dauti M, Wilde C, Mazel D, Rowe-Magnus DA. Chromosomal toxin-antitoxin loci can diminish large-scale genome reductions in the absence of selection. Molecular microbiology. 2007;63(6):1588–605. Epub 2007/03/21. doi: 10.1111/j.1365-2958.2007.05613.x. PubMed PMID: 17367382.

23. Biskri L, Mazel D. Erythromycin esterase gene *ere(A)* is located in a functional gene cassette in an unusual class 2 integron. Antimicrobial agents and chemotherapy. 2003;47(10):3326–31. Epub 2003/09/25. PubMed PMID: 14506050; PubMed Central PMCID: PMCPMC201170.

24. Stokes HW, Holmes AJ, Nield BS, Holley MP, Nevalainen KM, Mabbutt BC, et al. Gene cassette PCR: sequence-independent recovery of entire genes from environmental DNA. Applied and environmental microbiology. 2001;67(11):5240–6. Epub 2001/10/27. doi: 10.1128/aem.67.11.5240-5246.2001. PubMed PMID: 11679351; PubMed Central PMCID: PMCPMC93296.

25. Elsaied H, Stokes HW, Nakamura T, Kitamura K, Fuse H, Maruyama A. Novel and diverse integron integrase genes and integron-like gene cassettes are prevalent in deep-sea hydrothermal vents. Environmental microbiology. 2007;9(9):2298–312. Epub 2007/08/10. doi: 10.1111/j.1462-2920.2007.01344.x. PubMed PMID: 17686026.

26. Koenig JE, Sharp C, Dlutek M, Curtis B, Joss M, Boucher Y, et al. Integron Gene Cassettes and Degradation of Compounds Associated with Industrial Waste: The Case of the Sydney Tar Ponds. PloS one. 2009;4(4):e5276. doi: 10.1371/journal.pone.0005276.

27. Elsaied H, Stokes HW, Kitamura K, Kurusu Y, Kamagata Y, Maruyama A. Marine integrons containing novel integrase genes, attachment sites, *attI*, and associated gene cassettes in polluted sediments from Suez and Tokyo Bays. The ISME journal. 2011;5(7):1162–77. Epub 2011/01/21. doi: 10.1038/ismej.2010.208. PubMed PMID: 21248857; PubMed Central PMCID: PMCPMC3146285.

28. Tansirichaiya S, Rahman MA, Antepowicz A, Mullany P, Roberts AP. Detection of Novel Integrons in the Metagenome of Human Saliva. PloS one. 2016;11(6):e0157605. doi: 10.1371/journal.pone.0157605.

29. Chen X-L, Tang D-J, Jiang R-P, He Y-Q, Jiang B-L, Lu G-T, et al. sRNA-Xcc1, an integron-encoded transposon- and plasmid-transferred trans-acting sRNA, is under the positive control of the key virulence regulators HrpG and HrpX of *Xanthomonas campestris* pathovar *campestris*. RNA Biology. 2011;8(6):947–53. doi: 10.4161/rna.8.6.16690.

30. Coleman N, Tetu S, Wilson N, Holmes A. An unusual integron in *Treponema denticola*. Microbiology (Reading, England). 2004;150(Pt 11):3524–6. Epub 2004/11/06. doi: 10.1099/mic.0.27569-0. PubMed PMID: 15528643.

31. Brett PJ, Burtnick MN, Fenno JC, Gherardini FC. *Treponema denticola* TroR is a manganese- and iron-dependent transcriptional repressor. Molecular microbiology. 2008;70(2):396–409. Epub 2008/09/03. doi: 10.1111/j.1365-2958.2008.06418.x. PubMed PMID: 18761626; PubMed Central PMCID: PMCPMC2628430.

32. Limberger RJ, Slivienski LL, Izard J, Samsonoff WA. Insertional inactivation of *Treponema denticola tap1* results in a nonmotile mutant with elongated flagellar hooks. Journal of bacteriology. 1999;181(12):3743–50. Epub 1999/06/15. PubMed PMID: 10368149; PubMed Central PMCID: PMCPMC93852.

33. Paget MSB, Helmann JD. The σ^70^ family of sigma factors. Genome Biology. 2003;4(1):203–. PubMed PMID: PMC151288.

34. Koo B-M, Rhodius VA, Campbell EA, Gross CA. Mutational analysis of *Escherichia coli* σ^28^ and its target promoters reveal recognition of a composite −10 region, comprised of an “extended −10 motif” and a core-10 element. Molecular microbiology. 2009;72(4):830–43. doi: 10.1111/j.1365-2958.2009.06691.x. PubMed PMID: PMC2756079.

35. Stokes HW, O’Gorman DB, Recchia GD, Parsekhian M, Hall RM. Structure and function of 59-base element recombination sites associated with mobile gene cassettes. Molecular microbiology. 1997;26(4):731–45. doi: 10.1046/j.1365-2958.1997.6091980.x.

36. Boucher Y, Nesbø CL, Joss MJ, Robinson A, Mabbutt BC, Gillings MR, et al. Recovery and evolutionary analysis of complete integron gene cassette arrays from Vibrio. BMC evolutionary biology. 2006;6(1):3. doi: 10.1186/1471-2148-6-3.

37. Li X, Shi L, Yang W, Li L, Yamasaki S. New array of *aacA4-catB3-dfrA1* gene cassettes and a noncoding cassette from a class-1-integron-positive clinical strain of *Pseudomonas aeruginosa*. Antimicrobial agents and chemotherapy. 2006;50(6):2278–9. Epub 2006/05/26. doi: 10.1128/aac.01378-05. PubMed PMID: 16723608; PubMed Central PMCID: PMCPMC1479156.

38. Michael CA, Labbate M. Gene cassette transcription in a large integron-associated array. BMC genetics. 2010;11:82. Epub 2010/09/17. doi: 10.1186/1471-2156-11-82. PubMed PMID: 20843359; PubMed Central PMCID: PMCPMC2945992.

39. Papagiannitsis CC, Tzouvelekis LS, Miriagou V. Relative strengths of the class 1 integron promoter hybrid 2 and the combinations of strong and hybrid 1 with an active p2 promoter. Antimicrobial agents and chemotherapy. 2009;53(1):277–80. Epub 2008/11/13. doi: 10.1128/aac.00912-08. PubMed PMID: 19001114; PubMed Central PMCID: PMCPMC2612174.

40. Lévesque C, Brassard S, Lapointe J, Roy PH. Diversity and relative strength of tandem promoters for the antibiotic-resistance genes of several integron. Gene. 1994;142(1):49–54. doi: http://dx.doi.org/10.1016/0378-1119(94)90353-0.

41. Singh SS, Singh N, Bonocora RP, Fitzgerald DM, Wade JT, Grainger DC. Widespread suppression of intragenic transcription initiation by H-NS. Genes & development. 2014;28(3):214–9. Epub 2014/01/23. doi: 10.1101/gad.234336.113. PubMed PMID: 24449106; PubMed Central PMCID: PMCPMC3923964.

42. Jové T, Da Re S, Denis F, Mazel D, Ploy M-C. Inverse correlation between promoter strength and excision activity in class 1 integrons. PLOS Genetics. 2010;6(1):e1000793. doi: 10.1371/journal.pgen.1000793.

43. Guerin E, Jove T, Tabesse A, Mazel D, Ploy MC. High-level gene cassette transcription prevents integrase expression in class 1 integrons. Journal of bacteriology. 2011;193(20):5675–82. Epub 2011/08/23. doi: 10.1128/jb.05246-11. PubMed PMID: 21856858; PubMed Central PMCID: PMCPMC3187215.

44. Jacquier H, Zaoui C, Sanson-le Pors MJ, Mazel D, Bercot B. Translation regulation of integrons gene cassette expression by the *attC* sites. Molecular microbiology. 2009;72(6):1475–86. Epub 2009/06/03. doi: 10.1111/j.1365-2958.2009.06736.x. PubMed PMID: 19486293.

45. Collis CM, Hall RM. Expression of antibiotic resistance genes in the integrated cassettes of integrons. Antimicrobial agents and chemotherapy. 1995;39(1):155–62. Epub 1995/01/01. PubMed PMID: 7695299; PubMed Central PMCID: PMCPMC162502.

46. Guerout AM, Iqbal N, Mine N, Ducos-Galand M, Van Melderen L, Mazel D. Characterization of the *phd-doc* and *ccd* toxin-antitoxin cassettes from *Vibrio* superintegrons. Journal of bacteriology. 2013;195(10):2270–83. Epub 2013/03/12. doi: 10.1128/jb.01389-12. PubMed PMID: 23475970; PubMed Central PMCID: PMCPMC3650543.

47. Rowe-Magnus DA, Guerout AM, Biskri L, Bouige P, Mazel D. Comparative analysis of superintegrons: engineering extensive genetic diversity in the Vibrionaceae. Genome research. 2003;13(3):428–42. Epub 2003/03/06. doi: 10.1101/gr.617103. PubMed PMID: 12618374; PubMed Central PMCID: PMCPMC430272.

48. Van Melderen L, Saavedra De Bast M. Bacterial toxin-antitoxin systems: more than selfish entities? PLOS genetics. 2009;5(3):e1000437. Epub 2009/03/28. doi: 10.1371/journal.pgen.1000437. PubMed PMID: 19325885; PubMed Central PMCID: PMCPMC2654758.

49. Solovyev V, Salamov A. Automatic Annotation of Microbial Genomes and Metagenomic Sequences. In: Li RW, editor. In Metagenomics and its Applications in Agriculture, Biomedicine and Environmental Studies: Nova Science Publishers; 2011. p. 61–78.

50. Seier-Petersen MA, Jasni A, Aarestrup FM, Vigre H, Mullany P, Roberts AP, et al. Effect of subinhibitory concentrations of four commonly used biocides on the conjugative transfer of Tn916 in *Bacillus subtilis*. The Journal of antimicrobial chemotherapy. 2014;69(2):343–8. Epub 2013/10/05. doi: 10.1093/jac/dkt370. PubMed PMID: 24092655; PubMed Central PMCID: PMCPMC3886932.

51. Swartzman A, Kapoor S, Graham AF, Meighen EA. A new *Vibrio fischeri lux* gene precedes a bidirectional termination site for the *lux* operon. Journal of bacteriology. 1990;172(12):6797–802. Epub 1990/12/01. PubMed PMID: 2254256; PubMed Central PMCID: PMCPMC210795.

52. Dupuy B, Sonenshein AL. Regulated transcription of *Clostridium difficile* toxin genes. Molecular microbiology. 1998;27(1):107–20. doi: 10.1046/j.1365-2958.1998.00663.x.

53. Miller JH. Experiments in molecular genetics. NY: Cold Spring Harbor Laboratory Press: Cold Spring Harbor; 1972.

